# Pioneer factor Pax7 initiates two-step cell-cycle dependent chromatin opening

**DOI:** 10.1101/2022.11.16.516735

**Authors:** Arthur Gouhier, Justine Dumoulin-Gagnon, Vincent Lapointe-Roberge, Juliette Harris, Aurelio Balsalobre, Jacques Drouin

## Abstract

Pioneer transcription factors direct cell differentiation by deploying new enhancer repertoires through their unique ability to target and initiate remodelling of closed chromatin. The initial steps of their action remain undefined although pioneers were shown to interact with nucleosomal target DNA and with some chromatin remodelling complexes. We now define the sequence of events that enable the pioneer Pax7 with its unique abilities. Chromatin condensation exerted by linker histone H1 is the first constraint on Pax7 recruitment, and this establishes the initial speed of chromatin remodelling. The first step of pioneer action involves recruitment of the KDM1A (LSD1) H3K9me2 demethylase for removal of this repressive mark, as well as recruitment of the MLL complex for deposition of the activating H3K4me1 mark. Further progression of pioneer action requires passage through cell division, and this involves dissociation of pioneer targets from perinuclear lamin B. Only then, the SWI/SNF remodeling complex and the coactivator p300 are recruited, leading to nucleosome displacement and enhancer activation. Thus, the unique features of pioneer actions are those occurring in the lamin-associated compartment of the nucleus. This model is consistent with prior work that showed a dependence on cell division for establishment of new cell fates.

## Introduction

Cell fates are established and maintained through the action of specific combinations of transcription factors, including cell-restricted factors that define unique cell identities. The implementation of new cell fates relies on activation of new enhancer repertoires and this is achieved by pioneer transcription factors through their unique ability to access target sites in closed chromatin and trigger chromatin opening^1^. Many aspects of this general scheme remain undefined, notably pioneer interaction with closed chromatin and the initial events of chromatin remodelling^1^.

Investigation of DNA sequences targeted for pioneer action did not show preferential occurrence of sequence subsets, with some pioneers exhibiting degenerate recruitment sites^2^ while others have stronger recruitment at pioneered sites^3,4^. Closed chromatin access may include the ability to bind target DNA sequences on nucleosomes, one of the earliest features ascribed to many, but not all^5^, pioneer factors^6^. Structural evidence suggests that pioneer binding may weaken the interaction of DNA with nucleosomes and hence contribute to initial chromatin alteration^7–9^. This is compatible with the appearance of so-called “accessible nucleosome conformation” caused by FOXA recruitment^10^ or by subtle chromatin rearrangements that precede frank chromatin opening following the action of the pioneer Pax7 (REF.^3^). However, nucleosomal organisation cannot on its own explain barriers to pioneer action, and the data rather suggest that chromatin environment dictates whether chromatin is permissive or not for pioneer action. For example, the highly compacted constitutive heterochromatin prevents pioneer access^11,12^. The nature of crucial events initiating pioneer action remain elusive^1^.

It is likely that chromatin remodelling complexes identified for their role in development or in transcription are also involved in chromatin opening by pioneer factors. For example, the MLL complex is recruited to FOXA-pioneered sites that are targets of the oestrogen receptor in breast cancer cells^13^. The pioneer, Pax7, interacts with the methyltransferase CARM1 and the MLL1/2 proteins^14^. The pioneers OCT4, GATA3 and ISL1, interact with Brg1, the ATPase of the SWI/SNF chromatin-remodelling complex^15–17^. But it is presently unclear how and when these interactions may be relevant to pioneer-dependent chromatin opening.

In the present study, we used an inducible system to define the time course of chromatin opening at Pax7-pioneered enhancers^3^. Amongst its different roles^18^, Pax7 specifies the intermediate pituitary melanotrope cell fate by opening a repertoire of >2000 enhancers^3,19^. While Pax7 is necessary to trigger chromatin opening, it also requires cooperation with the nonpioneer transcription factor Tpit to fully open chromatin and activate enhancers ^20^. We now show that Pax7 pioneer action occurs in two steps, a first step that is limited by Pax7 recruitment strength and inversely correlated with the level of linker histone H1, and a second step that requires passage through replication and cell division for dissociation of large domains from lamin B, recruitment of the SWI/SNF complex, nucleosome displacement and enhancer activation. This sequential scheme of pioneer action defines a conceptual framework to further probe and control the pioneering process.

## Results

### Pax7 primes enhancers for activation

We previously provided an exhaustive analysis of chromatin opening by the pioneer factor Pax7 at two enhancer repertoires, one where Pax7 action results in complete enhancer activation and another where enhancers only reach the primed state^3^. To study the kinetics of enhancer opening by Pax7, we used a tamoxifen-inducible ER-Pax7 chimera system (Fig. 1a) that relies on expression of ER-Pax7 in AtT-20 cells at levels that are similar to those of mouse pituitary melanotrope cells (Extended Data Fig. 1a). This re-programs AtT-20 cells into melanotrope-like cells^3,19,20^. The present study compared subsets of enhancers that are either activated or primed in response to Pax7 with a set of constitutively active enhancers where Pax7 binding does not alter chromatin organization (Fig. 1b and Extended Data Fig. 1b, c). Having previously observed rapid Pax7 binding to pioneered sites in closed chromatin but relatively slower remodelling^3^, we first assessed whether the primed state is indeed a transitory state towards complete enhancer activation. The status of pioneered enhancers was assessed by ChIP-Seq for the activating chromatin mark histone monomethylated histone H3 Lys4 (H3K4me1) at different times following activation of ER-Pax7 (Fig. 1c). Thus, pioneered enhancers are observed first (12h) in the primed state followed by complete activation (Fig. 1c). The primed state was previously documented as presenting with a weak single peak of H3K4me1, whereas fully activated enhancers present with a stronger bimodal distribution of H3K4me1 that reflects nucleosome displacement at the center of pioneered enhancers^1^.

**Fig. 1.**
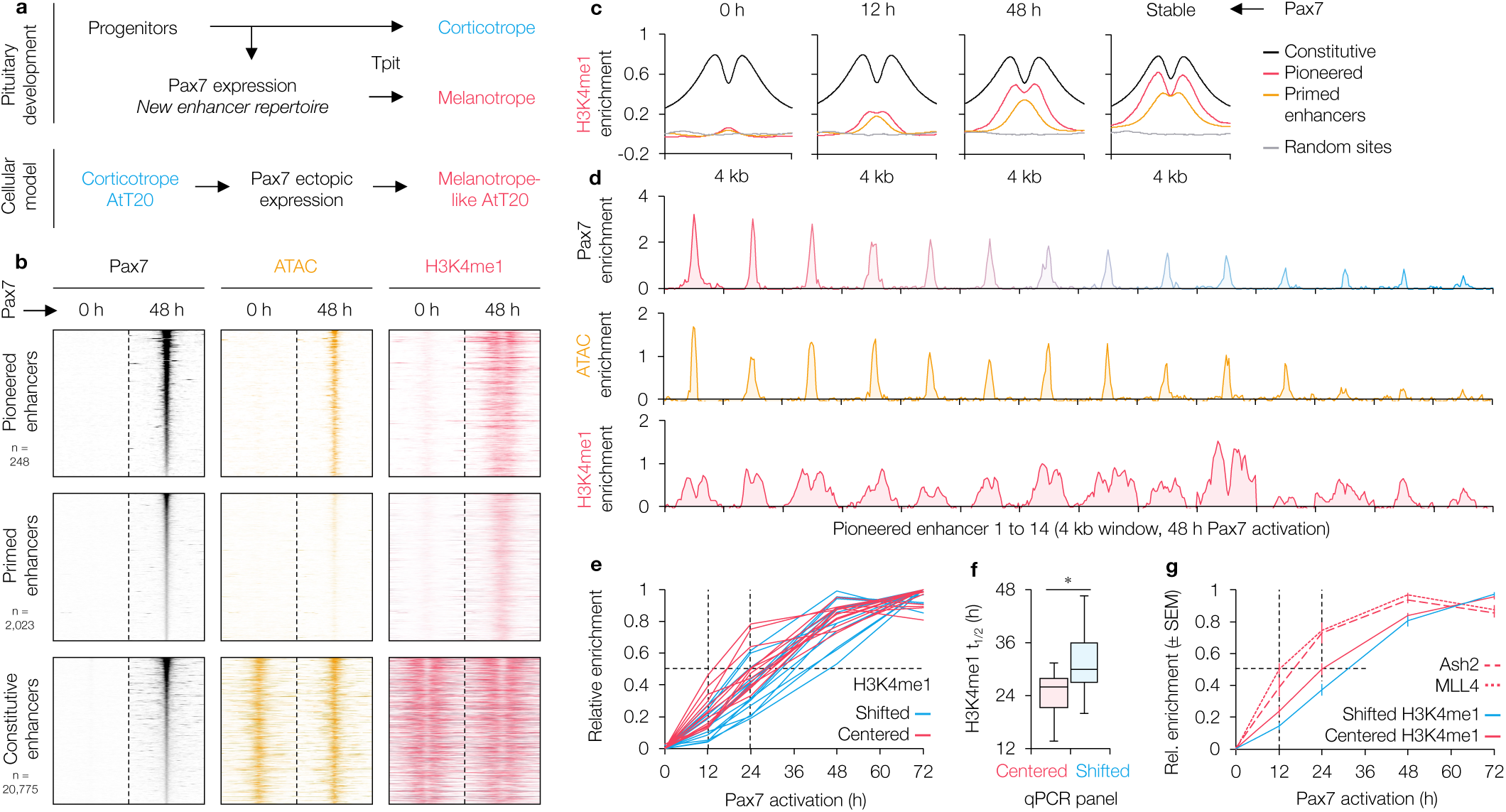
Pax7 primes enhancers for activation. **a,** The role of Pax7 in specification of the intermediate pituitary melanotrope cell fate is illustrated in contrast to the corticotrope cell fate. These two hormone-secreting lineages express the same hormone precursor gene, proopiomelanocortin (POMC), and their fates are determined by the Tbox transcription factor Tpit. The mouse AtT-20 cells are a model of the corticotrope lineage and Pax7 expression reprograms these cells towards a melanotrope-like fate. **b,** Heatmap representation of ChIP-Seq and ATAC-Seq data for three subsets of Pax7-targeted enhancers determined in Extended Fig. 1b before (0 h) and after 48 h of Pax7 activation (tamoxifen treatment) in AtT20-ERPax7 cells. Pioneered enhancers have no marks before Pax7 activation and present with hallmarks of transcriptionally active enhancers after, namely appearance of an ATAC-Seq signal together with a bimodal distribution of H3K4me1 deposition. Pax7 primed enhancers also show appearance of weaker ATAC-Seq and H3K4me1 signals, the latter presenting as a single weak peak. Finally, a set of constitutively active enhancers serve as control for comparisons. These enhancers bind Pax7, but this binding is not associated with any change in chromatin signature. These enhancer subsets were previously described^3,4^. **c,**) Mean profiles of H3K4me1 deposition at subsets of Pax7-remodelled enhancers at indicated times following tamoxifen (Tam) activation of ER-Pax7. **d,** Pax7 and H3K4me1 ChIP-Seq and ATAC-Seq signals at 14 randomly selected (Extended Data Fig. 1d) enhancers pioneered by Pax7 after 48 h of tamoxifen activation. Sites are sorted (red to blue) by the strength of Pax7 recruitment. **e,** Time course of H3K4me1 deposition determined by ChIP-qPCR at the 14 pioneered sites illustrated above. For each site, a red curve illustrates H3K4me1 accumulation under the Pax7 peak (centered), whereas a blue curve represents H3K4me1 deposition over a lateral peak of deposition (shifted) that characterizes the bimodal H3K4me1 distribution of fully active enhancers. **f,** Distribution of H3K4me1 deposition t_1/2_ from **(e)**. P-value from two-tailed t-test (*** ≤ 0.001, ** ≤ 0.01, * ≤ 0.5, – otherwise). **g,** Mean (± standard error of the mean, n ≥ 2) time course at 14 pioneered sites of H3K4me1 accumulation (centered and shifted) is compared with ChIP-qPCR data for recruitment of two protein components of the MLL remodelling complex, MLL4 and Ash2.

### Pioneered sites are heterogeneous

ChIP-Seq analyses of pioneered sites show significant heterogeneity with regards to Pax7 recruitment and deposition of chromatin marks (Fig. 1b). To assess the variability in implementation of Pax7 chromatin opening, we selected a panel of 14 pioneered sites chosen randomly and representative of the Pax7 recruitment distribution (Fig. 1d and Extended Data Fig. 1d). We validated that pioneered enhancers are first remodelled into a primed state by H3K4me1 ChIP-qPCR using two sets of primers, one centered on the Pax7 binding site and another shifted to one of the H3K4me1 bimodal deposition summit: indeed, H3K4me1 central deposition (average half-time of 24h) occurs before lateral (average half-time of 32h) accumulation (Fig. 1e,f and Extended Data Fig. 1e). Prior data had suggested interaction between Pax7 and the MLL complex that has H3K4me1 methyltransferase activity^14^. We therefore assessed MLL complex recruitment at Pax7 pioneered enhancers by ChIP-qPCR for the MLL3, MLL4 and Ash2 components of the MLL complex: both MLL4 and Ash2 are recruited with a half-time (average of the 14 sites) of ~12 hours following Pax7 activation (Fig. 1g and Extended Data Fig. 2). This half-time corresponds to a time when enhancers are in a primed state (Fig. 1c) and precedes bulk H3K4me1 deposition (Fig. 1f). This delay taken together with the prior demonstration of a delay between chromatin opening measured by ATAC-Seq and Pax7 recruitment^3^ suggests that chromatin opening initiated by Pax7 may involve sequential steps.

### Chromatin opening occurs in two steps

To assess the sequence of events leading to chromatin opening, we performed ChIP-qPCR time courses for various chromatin state markers and chromatin remodellers (Fig. 2a-d). Pax7 recruitment itself has an average half-time of ~12 hours and we found a similar time course for recruitment of the cooperating transcription factor Tpit (Fig. 2a). Recruitment of these factors parallels MLL4 and Ash2 recruitment, and they all precede deposition of H3K4me1 at the center of enhancers (Fig. 2a). We next assessed chromatin opening by ATAC-qPCR: the half-time of chromatin opening of ~24 hours is similar to H3K4me1 deposition (Fig. 2b). These longer time courses are similar to those of nucleosome displacement at these enhancers as revealed by ChIP-qPCR for histone H3 and for recruitment of Brg1 (Fig. 2b). The recruitment of the SWI/SNF complex (Brg1) occurs in parallel with recruitment of the general coactivator and histone acetylase p300 and of the cohesin complex protein SMC1 (Fig. 2b). These precede deposition of the active enhancer mark histone H3 acetylated Lys27, H3K27ac (Fig. 2c). These time courses clearly identify two steps in Pax7-dependent chromatin opening (Fig. 2a-c): an initial step marked by Pax7 recruitment and a delayed second step that involves nucleosome displacement (Fig. 2e, f).

**Fig. 2.**
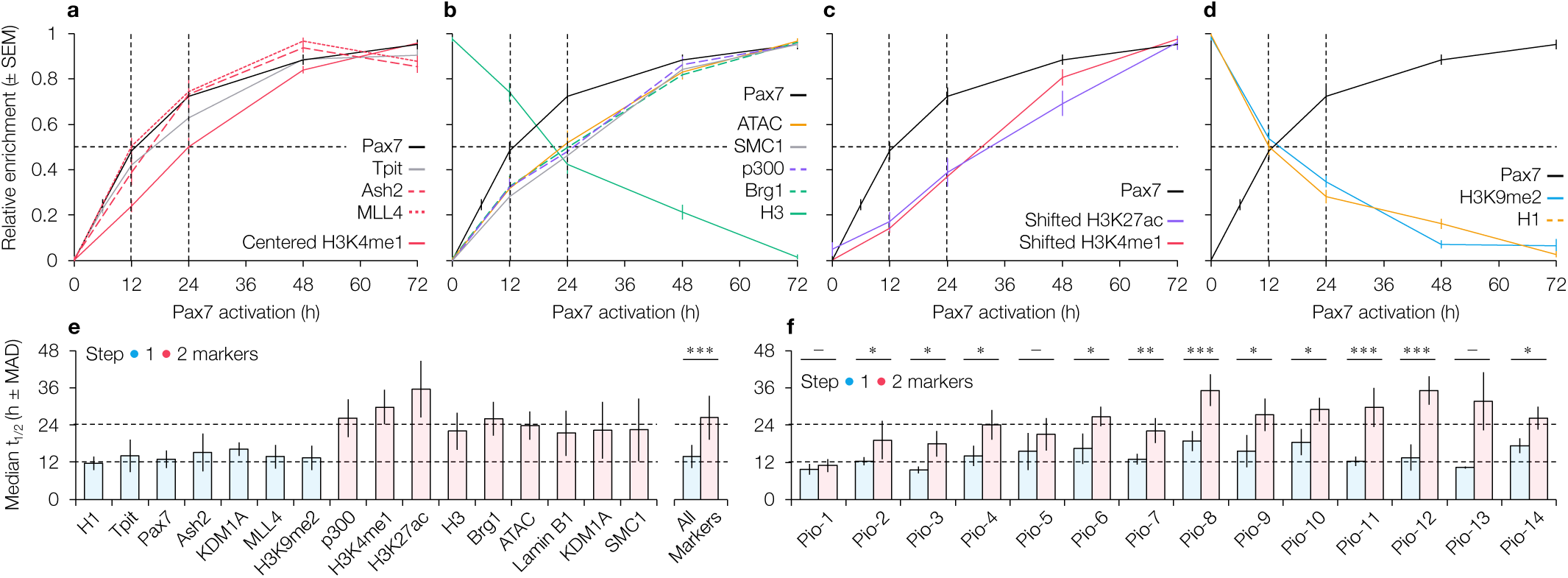
Pax7-dependent chromatin opening occurs in two steps. **a,** Average (14 pioneered sites of Fig. 1d represented as means ± sem of duplicate biological samples, n ≥ 2) profiles of ChIP-qPCR data for Pax7, Tpit, MLL4 and Ash2 that are recruited with a similar time course (T ½ ~ 12 hrs) compared to accumulation of H3K4me1 (center). **b,** Compared to Pax7 recruitment, ATAC-Seq signals and recruitment of Brg1, p300 and SMC1, together with depletion of Histone H3 occur with an average T ½ of ~ 24h (n ≥ 2). **c,** Deposition of activation mark H3K27ac and bimodal H3K4me1 (shifted) are slightly delayed compared to ATAC signal (n ≥ 2). **d,** Depletion of the repressive mark H3K9me2 and of linker Histone H1 occur with a T ½ of ~ 12h (n ≥ 2). **e,** Median (± median absolute deviation) half-times of change for the indicated parameter at the 14 pioneer sites. **f,** Median (± median absolute deviation) half-times for all parameters altered by Pax7 pioneering regrouped for step 1 (blue) and step 2 (pink) of the pioneering process. Data are shown for each of 14 pioneered enhancers (n ≥ 2). All panels: p-values from two-tailed t-tests (*** ≤ 0.001, ** ≤ 0.01, * ≤ 0.5, – otherwise).

We previously showed depletion of the repressive histone mark histone H3 dimethylated Lys9 (H3K9me2) that is typical of facultative heterochromatin at the Pax7 pioneered enhancers^3^. We assessed the time course of this depletion and observed a half-time of ~12 hours (Fig. 2d). Since linker histone H1 contributes to chromatin compaction, we measured its levels at the pioneered sites and observed a time course of H1 depletion similar to that of H3K9me2 (Fig. 2d). The initiation of Pax7 pioneering thus correlates with an initial perturbation of the chromatin environment represented by the loss of linker histone H1 and the loss of the repressive mark H3K9me2, together with implementation of the primed state reflected by a weak deposition of H3K4me1 (Fig. 1c).

### Pioneering kinetics depend on Pax7 recruitment strength

While the average time response curves defined two steps in the pioneering process, examination of parameters for the individual 14 pioneer sites revealed heterogeneity in the kinetics of pioneer action (Fig. 3a). Color-coded individual time course curves for Pax7 recruitment, histone H1 depletion, ATAC signal, H3K4me1 deposition, Brg1 recruitment and H3 depletion indicate that the stronger Pax7 recruitment sites (Fig. 1d, red) are remodelled quicker than the weaker sites (blue). The correlation between recruitment strength and half-time of remodelling is observed for all parameters (Fig. 3b). Similar correlations are observed for all pioneered enhancers genome-wide (Fig. 3c and Extended Data Fig. 3c). Computation of time course curves for the five strongest and five weakest sites indicates an average delay of ~6 hours between the two subgroups (Extended Data Fig. 3a, b). These data clearly support the conclusion that recruitment strength determines the onset of chromatin remodelling at pioneered sites.

**Fig. 3.**
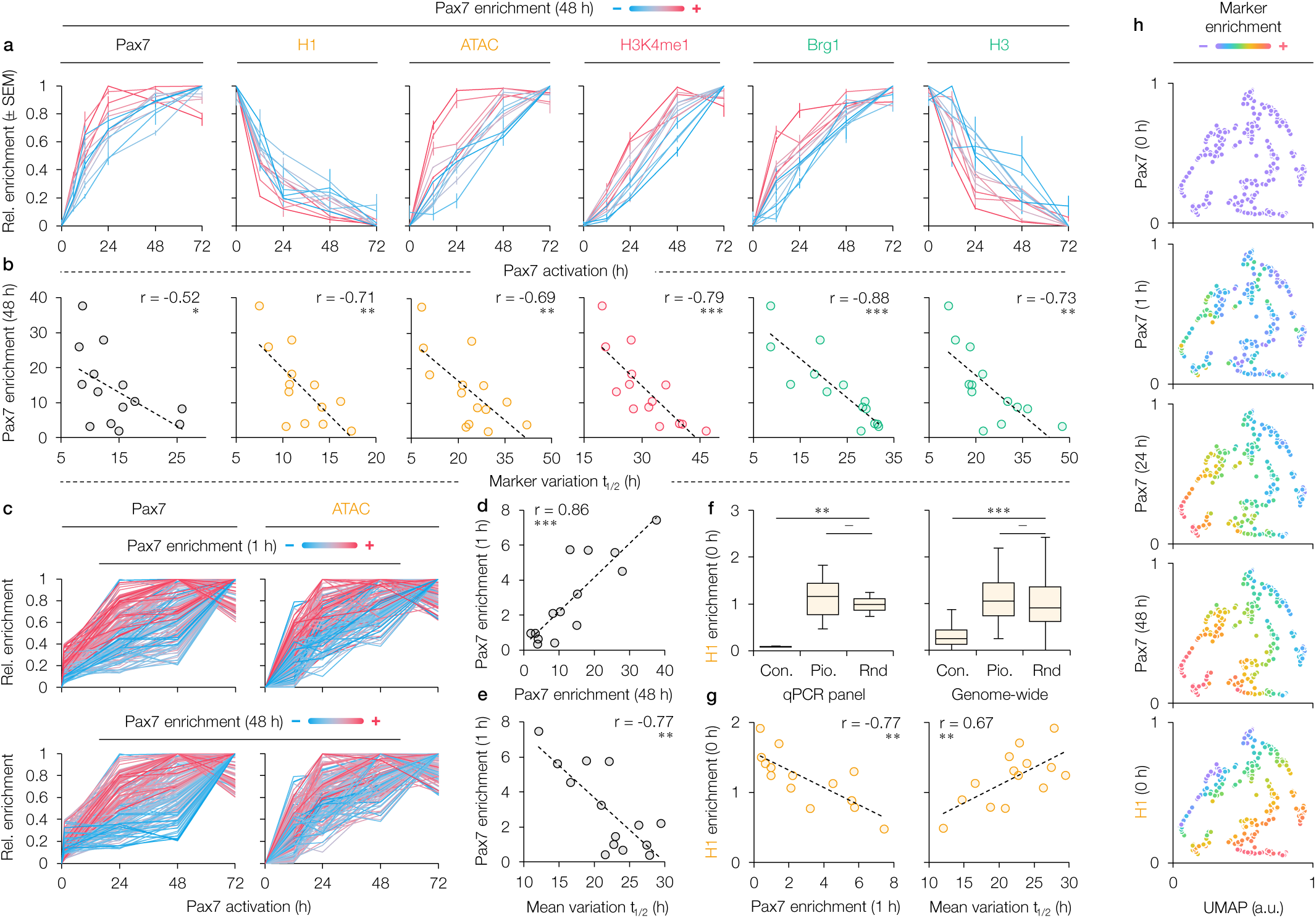
Initiation of Pax7 recruitment is delayed by histone H1-dependent chromatin compaction. **a,** Individual site ChIP-qPCR time course data (means ± standard error of the mean, n ≥ 2) for recruitment of Pax7, H3K4me1 and Brg1, together with time courses of ATAC signals and depletion histones H1 and H3 shown for the 14 pioneered sites color-coded (in Fig. 1d) from red to blue to reflect the strength of Pax7 recruitment. **b,** Correlations between Pax7 recruitment strength and various markers depicted in (a). **c,** Pax7 ChIP-Seq and ATAC-Seq time courses at all genome-wide pioneered enhancers. Sites color-coded according to Pax7 recruitment at 1 h and 48 h. **d,** Correlation at 14 pioneered enhancers between Pax7 recruitment signals at 1 h and 48 h. **e,** Inverse correlation between initial (1h) Pax7 recruitment strength and the average half-times of all markers of chromatin remodelling at the 14 pioneered sites. **f,** Distribution of initial histone H1 levels at constitutively active enhancers (con.), pioneered enhancers (pio.) and random sites (rnd) sites determined either genome-wide or from the qPCR panel (Extended Data Fig. 2). **g,** Correlations between initial histone H1 levels at the 14 pioneered enhancers and Pax7 recruitment at 1 h as well as with the mean t_1/2_ of all markers of the pioneering process. **h,** UMAP representation of Pax7 recruitment strengths at the indicated times after Tam addition compared to initial histone H1 levels measured by ChIP-Seq at pioneered enhancers depicted in Fig. 1b. All panels: p-values from two-tailed t-tests (*** ≤ 0.001, ** ≤ 0.01, * ≤ 0.5, – otherwise).

To assess whether these correlations relate to long-term or initial levels of Pax7 recruitment, we performed similar comparisons at 1 hour after Pax7 activation. These revealed that recruitment strength at 1 and 48 hours are directly correlated (Fig. 3d and Extended Data Fig. 4) and that initial recruitment is inversely correlated with the time course of chromatin remodelling (Fig. 3e).

We next assessed the impact on the strength of Pax7 recruitment of initial chromatin compaction as reflected by the levels of linker histone H1 that are higher at pioneered compared to constitutive sites (Fig. 3f). Whereas nucleosome content reflected by total histone H3 ChIP levels and the level of the repressive mark H3K9me2 (Extended Data Fig. 3d) do not correlate with Pax7 recruitment, chromatin compaction dependent on linker histone H1 inversely correlates with the initial (1h) strength of Pax7 recruitment (Fig. 3g). Similar observations were made at 48h after Pax7 activation (Extended Data Fig. 3c) and confirmed genome-wide by UMAP representation of Pax7 recruitment compared to histone H1 levels (Fig. 3h). Hence, the higher the level of histone H1, the weaker the initial Pax7 recruitment and the longer it takes to remodel these sites (Fig. 3h). It is noteworthy that long-term Pax7 recruitment and chromatin opening are not directly correlated with initial recruitment (Extended Data Fig. 4). In summary, chromatin condensation reflected by the level of histone H1 appear to constrain initial Pax7 recruitment and determine the time course of chromatin remodelling without affecting final pioneered enhancer activation.

### Activation of Pax7 primed enhancers requires cell division

The half-time of the second step of the pioneering process (~24 hours) corresponds roughly to the half-time for AtT-20 cell division. Thus, the second step of the pioneering process may require DNA replication or passage through cell division. We assessed this by blocking AtT-20 cell replication for 12 hours with mimosine (Fig. 4a, b). Mimosine-blocked AtT-20 cells are viable and can re-enter cell cycle upon release (Extended Data Fig. 5a). We assessed each step of the pioneering process by comparison of mimosine-blocked with normal cycling cells. These experiments indicate that Pax7 and Tpit recruitment, as well as H3K9me2 depletion, three parameters of the first pioneering step, occur independently of cell replication and division (Fig. 4c). In marked contrast, all marks of the second step of the pioneering process are blocked by mimosine treatment, including chromatin opening assessed by ATAC-qPCR, Brg1 and p300 recruitment, and deposition of H3K27ac (Fig. 4c and Extended Data Fig. 6). The mimosine treatment did not affect the levels of Brg1 and p300 but led to a ~2-fold decrease of Tpit (Extended Data Fig. 5b); however, this did not change Tpit recruitment in the first phase of Pax7 action (Fig. 4c). Since prior experiments showed that the first step of the pioneering process yields enhancers in the primed state, we assessed the pattern of H3K4me1 in mimosine-blocked cells that fail to undergo the second step of pioneering. The ChIP-Seq average profiles of pioneered enhancers in mimosine-blocked cells show that they remain in the primed state (Fig. 4d) in contrast to normal cycling cells in which they are in the activated state at a similar time following Pax7 activation (Fig. 1c). Also, mimosine-blocked cells only show minimal gains of ATAC-Seq signals (Fig. 4e) consistent with their primed status.

**Fig. 4.**
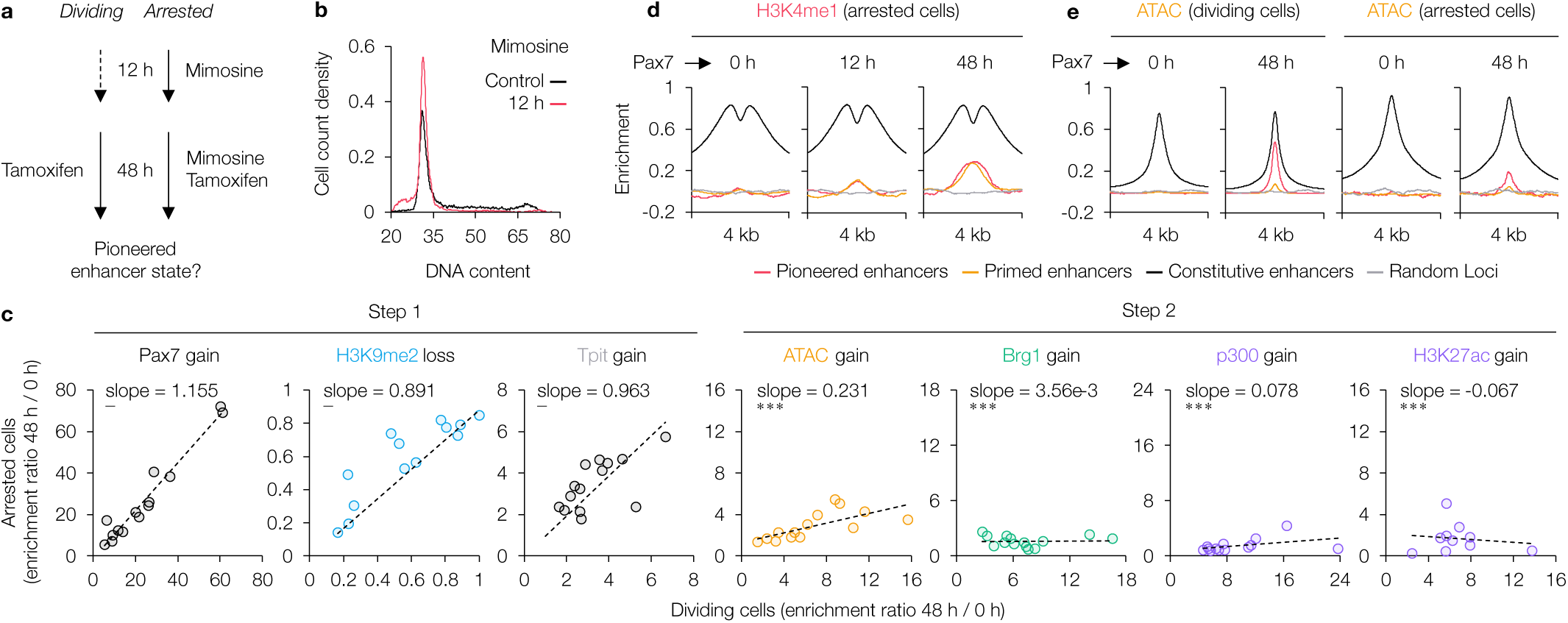
Activation of Pax7 pioneered enhancers requires cell division. **a,** Experimental design for assessment of chromatin remodelling in cells with mimosine-blocked cell cycle compared to normally cycling cells. **b,** FACS profiles of untreated cells compared to cells incubated for 12h with mimosine that exhibit cell-cycle arrest in G1. Representative of n=2 is shown. **c,** The recruitment of both Pax7 and Tpit, as well as depletion of H3K9me2, measured by ChIP-qPCR at the 14 pioneered sites are not affected by blocking AtT-20 cell cycle with mimosine (n=2). In contrast, DNA accessibility (ATAC-qPCR) as well as recruitments of Brg1, p300 and H3K27ac measured by ChIP-qPCR are all severely curtailed in mimosine-arrested cells (n≥2). P-values from two-tailed t-tests (*** ≤ 0.001, ** ≤ 0.01, * ≤ 0.5, – otherwise). **d, e,** Average profiles of H3K4me1 (**d**) and ATAC-Seq signals (**e**) accumulation at Pax7-remodelled enhancer subsets measured by ChIP-Seq at different times after Tam activation of ER-Pax7 in dividing or mimosine-arrested cells as indicated.

### Cell division is required for dissociation from nuclear lamins

Since the first change observed in parallel with Pax7 recruitment is the depletion of the repressive mark H3K9me2, we queried a putative mechanism for this depletion. The analysis of proteins associated with Pax7 in RIME experiments^21^ provided two candidate demethylases, KDM1A (LSD1) and KDM3A (Harris et al. In preparation). Hence, we assessed recruitment of these demethylases at Pax7 pioneered enhancers and found that KDM1A is recruited with a half-time corresponding to step 1 of the process (Fig. 5a) whereas KDM3A is recruited at step 2 (Extended Data Fig. 7a). The early loss of H3K9me2 may be a mechanism to alter the localization of pioneered sites in the nucleoplasm. Indeed, H3K9me2-marked loci are found in the nuclear periphery B compartment and their localization is altered by removal of H3K9me2 (REF.^22,23^). We thus assessed the association of the Pax7 pioneered sites with lamin B1, a marker of the nuclear B compartment (Fig. 5b) and found that high levels of lamin B1 correlated with high levels of H3K9me2 at pioneered sites (Fig. 5c, d). Despite the loss of H3K9me2 at step 1, we observed dissociation of pioneered sites from lamin B with a half-time corresponding to the cell cycle-dependent second step (Fig. 5a). If the requirement for cell division observed for step 2 of the pioneering process depends on nuclear compartment switching, the blockade of cell division should prevent dissociation from lamin B, and this is indeed what was observed in mimosine-blocked cells (Fig. 5e). In this context, we would expect that enhancers that are de novo primed by Pax7 would not show significant H3K9me2 depletion as is observed (Fig. 5f). We surmise that the slight decrease of H3K9me2 at primed sites is not sufficient for lamin B dissociation and this is supported by the similar levels of H3K9me2 at sites that were already in the primed state before Pax7 activation and that get transcriptionally activated by Pax7 (Activated enhancers, Fig. 5f). Further, the recruitment of the SWI/SNF complex is observed only after full activation but not with the primed state as revealed by Brg1 ChIP-Seq (Fig. 5f), in agreement with restricted nuclear localisation of Brg1 in the central nucleoplasm^24^. We found similar features for recruitment of RNA Pol II and components (MED1 and MED12) of the Mediator complex (Extended Data Fig. 7b). Why do some enhancers fail to complete the process of activation? A failure to recruit an essential cooperating transcription factor could be responsible and indeed, we observe no recruitment of Tpit, a nonpionner factor required for full activation of melanotrope enhancers^20^, at de novo primed sites (Fig. 5f).

**Fig. 5.**
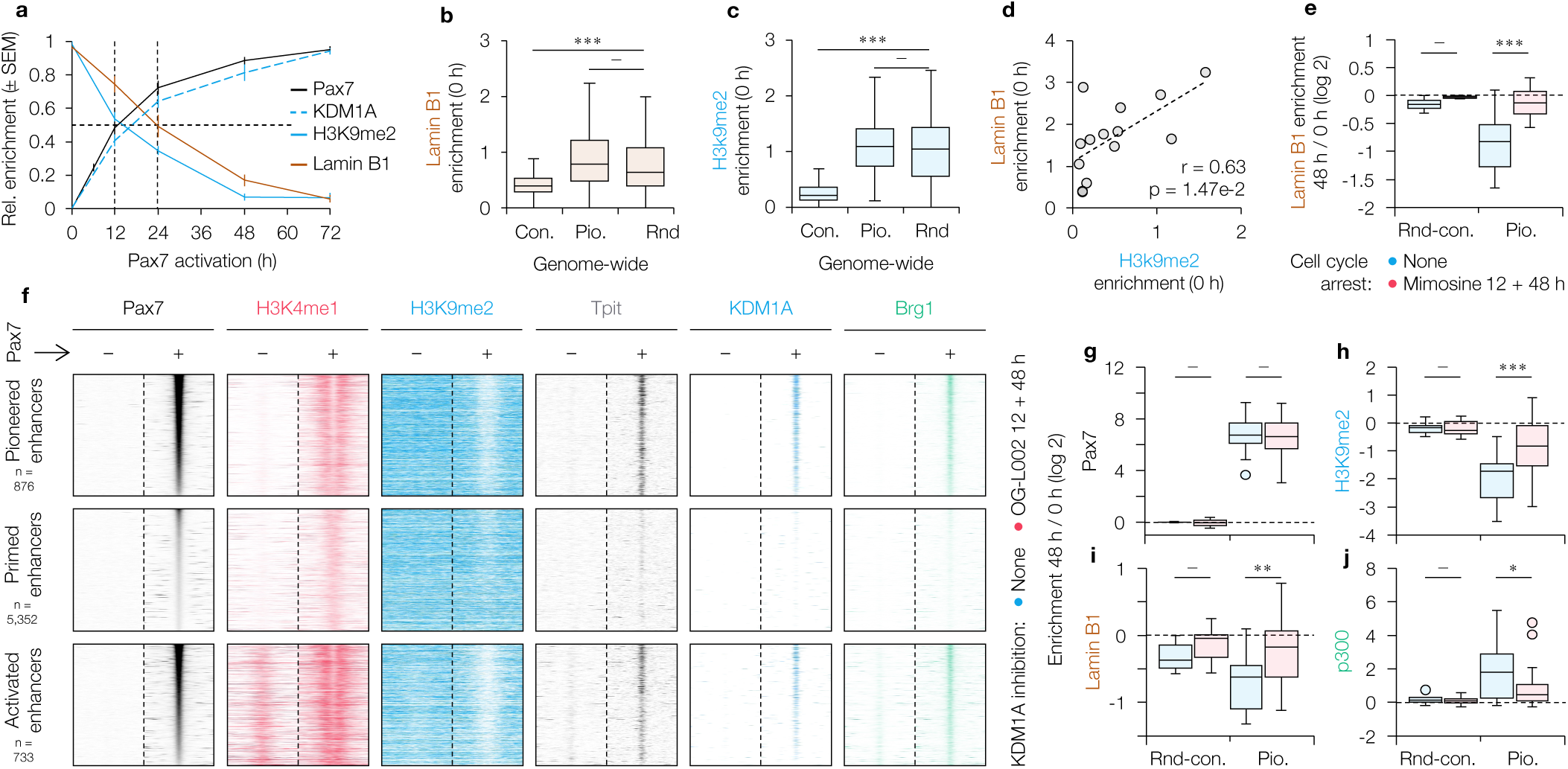
Pax7-dependent H3K9me2 demethylation and dissociation from lamin B. **a,** Mean (14 pioneered sites of Fig. 1d represented as means ± sem of duplicate biological samples) time courses for depletion of H3K9me2 and dissociation from lamin B1 measured by ChIP-qPCR compared to the time courses of recruitment for the H3K9me2 demethylase, KDM1A (LSD1). **b, c,** Pax7 pioneered sites have high levels (ChIP-Seq) of lamin B1 (**b**) and H3K9me2 (**c**) before Pax7 action. **d,** Correlation between initial levels (ChIP-qPCR, n ≥ 2) of H3K9me2 and lamin B1 at 14 pioneered sites. **e,** Ratios of Lamin B1 levels (ChIP-qPCR) at 48 over 0 h of Pax7 activation at constitutively active enhancers (con.), pioneered enhancers (pio.) and random loci (rnd) in dividing compared to mimosine-arrested AtT20 cells (n ≥ 2). **f,** Heatmaps of ChIP-Seq profiles for Pax7, H3K4me1, H3K9me2, Tpit, KDM1A(LSD1) and Brg1 at subsets of Pax7 remodelled enhancers. Data for other markers presented in Extended Data Fig. 7. **g-j,** ChIP-qPCR variation of Pax7 (g) and p300 (j) recruitment as well as H3K9me2 (h) and Lamin B1 (i) levels between no and 48 h of Pax7 activation at constitutively active enhancers (con.), the 14 pioneered enhancers (pio.) and random loci (rnd) in normal and cells treated (using highest concentration devoid of cytotoxicity) with the KDM1A inhibitor OG-L002 (n ≥ 2). Experimental design similar to mimosine treatment depicted in Fig. 4a. All panels: p-values from two-tailed t-tests (*** ≤ 0.001, ** ≤ 0.01, * ≤ 0.5, – otherwise).

In summary, the first step of pioneer action involves two biochemical alterations of critical histone marks: deposition of H3K4me1 presumably implemented by the MLL complex and removal of the repressive mark H3K9me2 by KDM1A. Both alterations occur independently of cell replication and division (Fig. 4c, d) but primed enhancers are characterized by the presence of the activation mark H3K4me1 without significant depletion of the repressive H3K9me2. We used an inhibitor of KDM1A to test whether its enzyme activity is essential for H3K9me2 demethylation and downstream chromatin changes. Whereas treatment with the KDM1A inhibitor OG-L002 does not prevent Pax7 recruitment (Fig. 5g), it decreases H3K9me2 demethylation (Fig. 5h), lamin B dissociation (Fig. 5i) and recruitment of p300 (Fig. 5j). These results support that KDM1A-dependent removal of the repressive H3K9me2 mark is needed for lamin B dissociation and complete enhancer activation. Further, the pioneer-dependent priming of enhancers only depends on H3K4me1 deposition independently of H3K9me2 depletion.

### TAD size lamin B dissociation initiated by Pax7

We next assessed the extent of epigenetic changes associated with pioneer-dependent chromatin opening using qPCR ChIP measurements of histones H3 and H1, together with levels of lamin B association at three pioneered loci (Fig. 6). These loci are enriched for H3K9me2 and lamin B (Fig. 6a, b) and ATAC-Seq profiles performed on wild-type and Pax7 mutant mouse pituitary tissues defined TAD size domains (Fig. 6a) that show Pax7-dependent opening (Fig. 6b). In contrast to the highly localized nucleosome depletion revealed by histone H3 levels at the center of Pax7 pioneered enhancers, displacement of histone H1 is more variable, appears locus-specific and may extend over several Kb’s (Fig. 6c, d). The dissociation from lamin B is more striking as it extends the full length of TADs containing the pioneered sites (Fig. 6). This is consistent with our prior observation of TAD size chromatin opening at loci containing melanotrope-specific genes such as *Pcsk2* and *Drd2* ^20^.

**Fig. 6.**
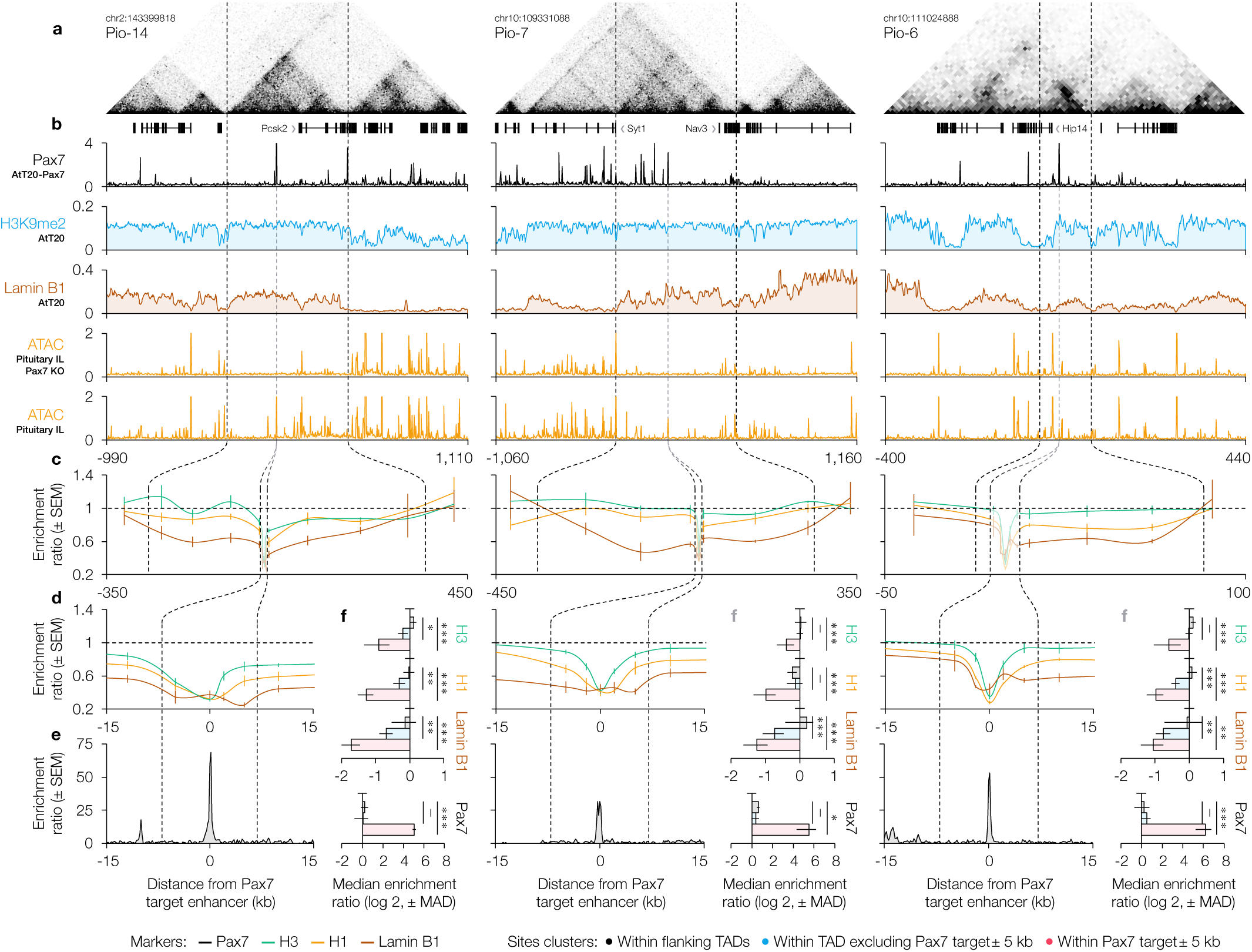
Pax7 initiates domain-wide lamin B dissociation. **a,** HiC contact scores in neural cells^37^ and annotated genes at three TADs harboring a Pax7-pioneered enhancer (including the hallmark Psck2 enhancer, Pio-14) and their flanking regions. **b,** Stable Pax7 ChIP-seq recruitment in AtT20-Pax7 cells. H3K9me2 and Lamin B1 ChIP-seq in AtT20 cells. ATAC-seq in WT and Pax7-KO intermediate lobe (IL) mouse pituitary cells depicting the dependance on Pax7 for activation of these TADs. Units: reads per million. **c,** Mean changes (± standard error of the mean, n ≥ 3) in histones H3, H1 and Lamin B1 levels following stable Pax7 expression in AtT20 cells. Data acquired by ChIP-qPCR at numerous sites across target TADs and flanking regions. By contrast to the local histone H3 and H1 depletions, Lamin B1 dissociation spans the entirety of the TADs. Flanking regions are not affected by Pax7. **d,** Same as **(c)** but zoomed on Pax7-targeted enhancer. **e,** Local Pax7 recruitment by ChIP-Seq at the pioneered enhancers. **f,** Distribution of Pax7 recruitment, histone H3 and H1 and Lamin B1 changes depicted in **(c)** with measured sites clustered as: flanking TADs, within target TAD excluding Pax7 target ± 5 kb and within Pax7 target ± 5 kb. P-values from two-tailed t-tests (*** ≤ 0.001, ** ≤ 0.01, * ≤ 0.5, – otherwise).

Whereas the three loci have different numbers of Pax7 recruitment sites and/or Pax7-dependent changes in ATAC-Seq profiles, the depletion of histones H3 and H1 is localized around the Pax7 peaks in contrast with lamin B dissociation. These three loci have only one (Pio-14 and -6) or two (Pio7) pioneered enhancers; the other Pax7 recruitment sites were in the primed state before Pax7. By contrast to Pio7 that shows many Pax7 recruitment sites, loci Pio14 and Pio6 only have one other significant Pax7 sites that may contribute to lamin B dissociation. It thus appears that a single pioneered site may be sufficient to drive domain-scale lamin B dissociation.

In conclusion, and taken collectively, the present data suggest a model for pioneer factor action (Fig. 7) in which the first steps are compatible with actions within the perinuclear compartment B: these include removal of repressor marks together with deposition of activation marks corresponding to the primed state. The ensuing steps for enhancer opening require passage through cell division that allows for lamin dissociation, and presumably switching to the A compartment, where the loci become accessible to the SWI/SNF complex for nucleosome displacement and transcriptional activation.

**Fig. 7.**
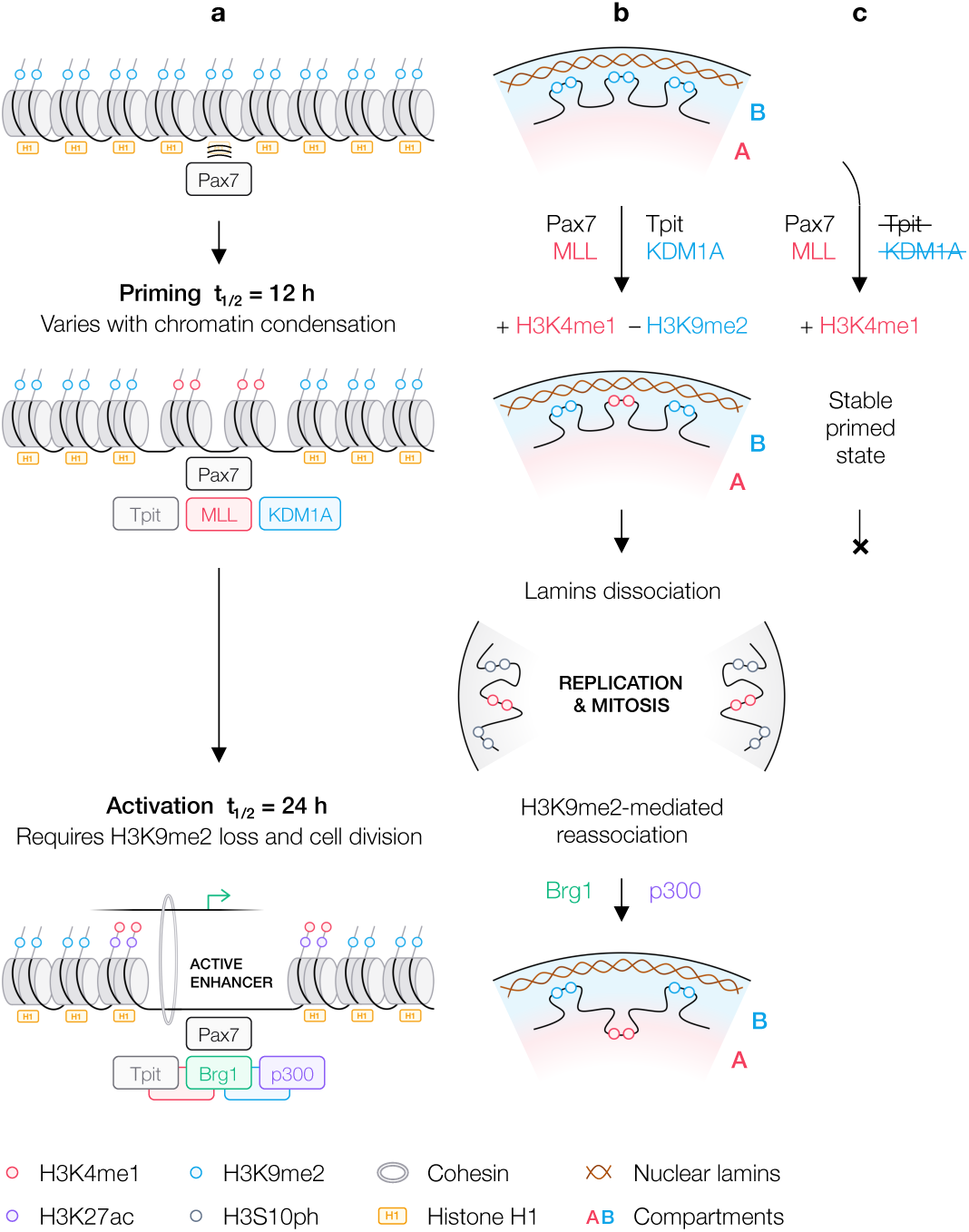
Model of Pax7-initiated chromatin opening and enhancer activation. **a,** Schematic representation of proteins shown in the present work to be associated with either primed or active enhancer states. **b,** Sequential model of Pax7 pioneer action. Pax7 pioneer targets are enriched in H3K9me2 and associated with lamin B1, which is typical of chromatin that is present in the perinuclear B compartment. Step 1 of the pioneering process involves recruitment of Pax7 and Tpit together with the H3K9me2 demethylase KDM1A (LSD1) and the MLL complex. These lead to demethylation of H3K9me2 and H3K4me1 deposition, respectively. Cell cycle blockade with mimosine prevents Pax7-dependent remodelling from proceeding further. At mitosis, masking of the H3K9me2 mark by H3S10 phosphorylation accompanies dissociation of A and B compartments of the nucleus^22^. Following mitosis and nuclear reassembly, Pax7-pioneered targets that have undergone H3K9me2 demethylation will remain in the central nuclear A compartment where the Brg1 and Brm ATPase’s of the SWI/SNF complex are localised^24^ and recruited for nucleosome displacement and complete enhancer activation. **c,** Pax7 primed enhancers do not recruit the cooperating nonpioneer factor Tpit and hence, the cooperation between Tpit and Pax7 appears required for significant H3K9me2 demethylation and continuation of the Pax7-dependent remodelling process to the cell cycle-dependent step 2.

## Discussion

The present work shows that chromatin compaction elicited by histone H1 determines the initial strength of pioneer Pax7 recruitment and speed of its actions. Beyond this limiting step, the different parameters of chromatin opening follow a similar time course while the next limiting step is imposed by cell division and its importance for lamin B dissociation (Fig. 7). Lamin B association has been related to the levels of H3K9 methylation and the strength of this association correlated with localization within perinuclear compartment B^22,23^ or with subdomains of this compartment^25^. Whereas the extent of lamin B association is correlated with the level of H3K9me2 at various loci (Fig. 5d), demethylation of H3K9me2 occurs earlier (step1) than lamin dissociation (step 2) with the latter coinciding with cell division (Fig. 5a). Although pioneered loci were not visualized within the nucleus in the present work, their association with lamin B suggest that the Pax7 pioneer target enhancers are localized within the perinucleoplasmic compartment B. The association of large LAD domains with the nuclear lamina and their dissociation following the loss of the H3K9me2 mark^22,23^ is consistent with our qPCR-ChIP data on lamin B association of pioneered loci and the TAD-size decrease of lamin B association following Pax7-dependent remodeling (Fig. 6). This large scale lamin B dissociation contrasts with the more localized depletion of histone H1 around Pax7 pioneered enhancers that also exhibit nucleosome displacement. The relationship between localized Pax7-dependent enhancer chromatin opening and the TAD size rearrangements remain to be elucidated. Whether the decreased lamin B association of pioneered enhancers reflects compartment (or sub-compartment^25^) switching or not, the apparent dissociation reflects a significant change of organization as it is correlated with the ability for full activation including recruitment of the SWI/SNF ATPase Brg1 that is only found in nucleoplasmic compartment A ^24^.

Both our previous analyses^3^ and recent papers^26,27^ showed that pioneered sites exhibit strong pioneer recruitment whereas weaker sites are more frequently associated with the primed states although there is significant overlap between these subsets. These observations led some authors to suggest that recruitment strength is critical for pioneer action^26,27^. We found that Pax7 isoforms or mutants that have decreased affinity fail to open subsets of targets but these analyses also revealed intrinsic target site differences that presumably reflects context-dependent chromatin heterogeneity^4,28^. Irrespective of these relations, the present work suggests a model where permissive sites of Pax7 pioneer action are in various states of compaction as reflected by the level of histone H1. While this compaction appears to hamper initiation of Pax7 action, it does not prevent it as long-term levels of Pax7 recruitment and action are not directly related to the 1h and 48h levels (Extended Data Fig. 4). Hence, the impact of histone H1 compaction is limited to initial Pax7 recruitment.

In parallel with histone H1 displacement, Pax7 recruits the MLL complex that has histone H3K4me1 methylase activity: enhancers that are primed after Pax7 recruitment have this signature (Fig. 7c). The priming of enhancers is thus possible without dissociation from lamin B (and possibly compartment B). Complete enhancer opening by Pax7 involves recruitment of the cooperating factor Tpit and of the H3K9me2 demethylase KDM1A (Fig. 7b): the activity of this enzyme is, at least partly, responsible for H3K9me2 demethylation and for subsequent steps of enhancer activation, including dissociation from lamin B (Fig. 5g-j). It will be interesting to compare the stepwise process described here for Pax7-dependent pioneer action with other pioneers. While there is as yet no similar temporal description of pioneer action, a two-tier process that is consistent with our model was proposed for Zelda^29^. The targets of many pioneers, or the pioneers themselves, were shown to recruit the MLL^13,14^ and SWI/SNF^5,15,30^ complexes; however, their sequence of action or their relative involvement at the naïve or primed states are not documented. Similarly, KDM1A was associated with the actions of pioneers, but also many TFs^31,32^.

The realization that Pax7 pioneering targets are enriched in H3K9me2 and that removal of this mark by KDM1A is required for activation raises the question of the nature of permissive sites for other pioneers. While it is quite possible that different pioneers may target different “closed” chromatin, it is difficult to assess the H3K9me2 status of chromatin sites targeted by other pioneers as this mark was either not investigated or the naïve versus primed status of the targets not sorted out^31^. Irrespective of pioneer involvement, the association of differentiation genes with LAD switching from compartments B to A is however documented in many systems^25,33^.

Similar limitations make it difficult to compare the role of replication or cell division in pioneer action. This was investigated for FOXA-mediated chromatin opening by *Donaghey* et al.^34^ who showed that blockade of replication prevents FOXA-dependent DNA demethylation without affecting bulk chromatin opening. Meaningful comparisons between different studies would require clear distinction of pioneer target subsets as either naïve or primed because this impacts dependence on replication for enhancer activation. Most cell reprogramming systems^35,36^ typically take place over a few days and would involve cell divisions. As far as we can tell, the link between pioneering and replication is indirect as cell division appears required for nuclear compartment switching^22^ rather than for the unique aspects of pioneer action, namely the priming that results in H3K4me1 deposition (Fig. 7). The recruitment of KDM1A to the pioneer Pax7 may thus represent a mechanism for chromatin opening at enhancers that are within H3K9me2-associated LADs. Whether recruitment of the MLL complex and of KDM1A are due to direct interactions with Pax7 or Tpit respectively, or whether they are interdependent, remains to be established. But the independence of the two critical changes of chromatin marks at pioneered enhancers supports models where a single pioneer factor may implement various subsets of primed enhancers ready for activation in response to an array of developmental (such as Tpit in our system) or signaling (such as Stat3 and GR^3^) cues mediated through the activities of different nonpioneers.

## ACKNOWLEDGMENTS

We are very grateful to Dr. Ali Shilatifard for the MLL3 and MLL4 antibodies, to Odile Neyret and Sarah Boisset for NGS analyses, and to Valérie Magoon for her expert secretarial assistance. Data analyses were possible thanks to the support of Compute Canada. This work was supported by Foundation grant FDN-154297 to J.D. from the Canadian Institutes of Health Research.

## AUTHOR CONTRIBUTIONS

A.G. and J.D. conceived the study, A.G., A.B. and J.D. conceived and designed the experiments, A.G., J.D.-G., V.L.-R. and J. H. performed experiments, A.G., V.L.-R. and J.D. wrote the manuscript.

