## Extended Data Figures and Methods for "Pioneer factor Pax7 initiates two-step cell-cycle dependent chromatin opening"

### Extended Data Information

### Methods

**Cell culture.** AtT20 cells (obtained from the late E. Herbert in 1981 and subsequently maintained in our laboratory, with yearly negative mycoplasma tests) were cultured in Dulbecco's modified Eagle's medium (DMEM) supplemented with 10 % fetal bovine serum (FBS) and antibiotics (penicillin, streptomycin). AtT20-Pax7 and AtT20-ERPax7 transgenic cells were previously described[3,19]. Tamoxifen (Sigma-Aldrich H7904) treatment was at a final concentration of 400 nM and 0.1 % ethanol. Mimosine (Sigma-Aldrich M0253) treatment was at a final concentration of 200 µM and OG-L002 (Tocris Bioscience 6244) at a final concentration of 50 µM. For treatments exceeding 24 h, media were replaced w/wo indicated agents every day.

**Cell cycle analysis.** Cells were fixed in 70 % ethanol, washed with PBS and incubated with 100 µg/ml RNase and 15 µg/ml propidium iodide for 30 min. Propidium iodide fluorescence analyses to assess DNA content per cell were performed on FACSCalibur cell analyzer using CellQuest Pro (BD Bioscience) on ~10,000 cells.

**Cell extracts.** Cells were incubated 15 min on ice in low salt buffer (10 mM HEPES pH 7.9, 10 mM KCl, 0.1 mM EDTA, 0.1 mM EGTA) before NP40 was added to a final concentration of 1 %. The cytoplasmic fraction was extracted and nuclear fraction incubated 30 min at 4 °C in high salt buffer (20 mM HEPES pH 7.9, 400 mM KCl, 1 mM EDTA, 1 mM EGTA). Protein concentrations were quantified by Bradford assay.

**Chromatin immunoprecipitation.** ChIP were performed as previously described[38] . Briefly, cells were crosslinked 10 min at room temperature or 37 °C in 1 % formaldehyde and incubated 5 min in quenching buffer (125 mM glycine in PBS). Cells were lysed by 5 min and 30 min incubation on ice in lysis buffers (10 mM tris-HCl pH 8, 0.25 % Triton X-100, 10 mM EDTA, 0.5 mM EGTA followed by 10 mM tris-HCl pH 8, 200 mM NaCl, 1 mM EDTA, 0.5 mM EGTA). Sonication was performed using Covaris E220 in sonication buffer (10 mM tris-HCl pH 8, 140 mM NaCl, 0.5 % Triton X-100, 0.5 % SDS, 0.05 % NaDOC, 1 mM EDTA, 0.5 mM EGTA or 20 mM tris-HCl pH 8, 150 mM NaCl, 0.1 % SDS, 1 % Triton X-100, 2 mM EDTA) to maximize DNA fragments of 100 to 500 bp. Antibody-Protein A/G coupled magnetic beads (Cytiva 17152104010150) were incubated overnight at 4 °C with input chromatin from 10 million cells and washed by serial incubation in wash buffers (20 mM tris-HCl pH 8, 150 mM NaCl, 0.1 % SDS, 1 % Triton X-100, 2 mM EDTA; 20 mM tris-HCl pH 8, 500 mM NaCl, 0.1 % SDS, 1 % Triton X-100, 2 mM EDTA; 10 mM tris-HCl pH 8, 250 mM LiCl, 1 % NP40, 1 mM EDTA; 10 mM tris-HCl pH 8, 50 mM NaCl, 1 mM EDTA). Crosslink was reversed by overnight incubation at 65

°C in elution buffer (50 mM tris-HCl pH 8, 1 % SDS, 2 mM EDTA) and DNA fragments purified following 1 h incubation at 65 °C with RNase and proteinase K. Fragments for H1 and Lamin B1 ChIP-Seq were digested further to 100 to 500 bp by 12 min incubation at 37 °C with 5.7 U/mg DNase I in digestion buffer (10 mM mM tris-HCl pH 8, 2.5 mM MgCl2, 0.5 mM CaCl2) and the reaction stopped by 10 min incubation at 75 °C with 5 mM EDTA.

**Assay for transposase-accessible chromatin.** ATAC were performed as previously described[3]. Briefly, nuclei from 50,000 cells were isolated by 30 min incubations on ice in lysis buffers (0.1 % sodium citrate tribasic dihydrate, 0.1 % Triton X-100 followed by 10 mM tris-HCl pH 7.4, 10 mM NaCl, 3 mM MgCl2, 0.1 % IGEPAL CA-630). Transposition was performed on nuclei using Tagment DNA kit following Illumina's recommendations and DNA fragments purified.

**Quantitative PCR analyses.** qPCR was performed as previously described[4] using SYBR Green (Applied Biosystems A25741) on ViiA-7 (Applied Biosystems). Signal by locus was normalized to fragmented input DNA (standard curve) and expressed relatively to average signal at random loci and/or constitutively active enhancers. Loci and primers are listed in Supplementary Table 1.

**Genome-wide analyses.** Libraries and flow cells were prepared by the IRCM Molecular Biology Core Facility according to Illumina's recommendations and sequenced on Illumina HiSeq, NextSeq or NovaSeq sequencers. Sequencing parameters are listed in Supplementary Table 2. Paired-end reads were first trimmed for adapter content and aligned to the mouse mm10 reference genome using Bowtie2/2.4 (--no-unal --no-mixed --no-discordant)[39]. Bam files were created and duplicated reads removed using view (-b) and markdup (-r) from SAMtools/1.12 (REF. [40]). Reads from multiple biological replicates were merged (if applicable). Coverage tracks were created using bamCoverage (-bs 10 -e 150 --normalizeUsing BPM) from deepTools/3.5 (REF. [41]). Peaks were called using callpeak (-f BAMPE -p 1e-3/1e-5 -g mm --min-length 100 --max-gap 50) from MACS/2.2.7.1 (REF. [42]). Local signal was quantified by the Euclidean norm over a 1,000 bp window (2,000 bp for H3K4me1 and H3K27ac). Pax7-bound loci localized within a 1,000 bp window of a gene's TSS (GENCODE M25) or with an input signal greater than the 3$^{rd}$ quartile + 3 * IQR of all bound loci's signal were filtered out. Pax7-bound loci were categorized as previously defined[4] for the stable system and as shown in Extended Data Fig. 1a for the inducible system. ChIP-Seq and ATAC-Seq signals were normalized relative to the average signals at constitutively active enhancers. DNA binding motifs were scored using annotatePeaks from Homer/4.11 (REF. [43]) over a 100 bp window.

**Time course analyses.** For each chromatin marker, normalized ChIP and ATAC signal per locus was relativized to the total variation between 0 and 72 h of Pax7 activation. Aberrant locus time course replicates were filtered out if the sum of incorrect variation (i.e., in the opposite direction of the total variation) between two time-points exceeded 40 % of the total variation or if any variation between two time-points exceeded 80 % of the total variation. For qPCR data, at least two biological replicates per experimental condition were merged.

**Antibodies.** Flag (FLAG M2): Sigma-Aldrich F3165. Pax7 (PAX7): DSHB AB_528428 for WB, Flag antibody for ChIP. Tpit (TPIT): Drouin lab, IRCM. MLL3 (KMT2C) and MLL4 (KMT2D): Shilatifard lab, Northwestern University. Ash2 (ASH2L): Bethyl A300-489A. Brg1 (SMARCA4): Abcam ab110641. p300 (EP300): Abcam ab14984. Lsd1 (KDM1A): Abcam ab17721. JmjD1 (KDM3A): Abcam ab243641. Pol II (POLR2A): Santa Cruz sc-899. Med1 (MED1): Bethyl A300-793A. Med12 (MED12): Bethyl A300-774A. SMC1: Bethyl A300-055A. Lamin B1 (LMNB1): Abcam ab16048. H1 (H1.4, H1.5): Millipore 05-457. H3: Abcam ab1791. H3K4me1: Abcam ab8895. H3K27ac: Abcam ab4729. H3K9me2: Abcam ab1220. H3K9me3: Millipore 07-442.

### References.


38 Langlais, D., Couture, C. & Drouin, J. The Stat3/GR interaction code: predictive value of direct/indirect DNA recruitment for transcription outcome. *Molecular Cell* **47**, 38-49 (2012).

39 Langmead, B., Trapnell, C., Pop, M. & Salzberg, S. L. Ultrafast and memory-efficient alignment of short DNA sequences to the human genome. *Genome Biol* **10**, R25 (2009).

40 Li, H. *et al.* The Sequence Alignment/Map format and SAMtools. *Bioinformatics* **25**, 2078-2079 (2009). https://doi.org:10.1093/bioinformatics/btp352

41 Ramírez, F. *et al.* deepTools2: a next generation web server for deep-sequencing data analysis. *Nucleic Acids Res* **44**, W160-165 (2016). https://doi.org:10.1093/nar/gkw257

42 Zhang, Y. *et al.* Model-based analysis of ChIP-Seq (MACS). *Genome Biol* **9**, R137 (2008). https://doi.org:10.1186/gb-2008-9-9-r137

43 Heinz, S. *et al.* Simple combinations of lineage-determining transcription factors prime cis-regulatory elements required for macrophage and B cell identities. *Mol Cell* **38**, 576-589 (2010).

**Extended Data Fig. 1 | Genome-wide effect of Pax7. a,** Pax7 western blot of total cell extracts from neurointermediate lobe (IL) pituitary cells (~50 % melanotropes), AtT20 cells and AtT20-ERPax7 cells showing ectopic expression of ERPax7 at physiological levels (n = 3). **b,** H3K4me1 ChIP-Seq and ATAC-Seq signals at Pax7 binding sites before and after 48 h of Pax7 activation. Bottom plots show selection threshold for constitutively active, pioneered and primed enhancers bound by Pax7. **c,** Boxplots of Pax7 ChIP-Seq, H3K4me1 ChIP-Seq and ATAC-Seq signals at random, constitutively active and pioneered sites as defined in **(b)** from no to 72 h of Pax7 activation and with stable Pax7 expression. **d,** Frequency per bin of Pax7 ChIP-Seq signals at pioneered enhancers binned according to signal strength. The randomly selected 14 pioneered enhancers used in the qPCR panel are indicated by vertical bars (red lines). **e,** H3K4me1 ChIP-qPCR variation $t_{1/2}$ at the 14 pioneered enhancers measured either at the Pax7 binding site (centered) or shifted towards one of the lateral H3K4me1 deposition summits.

**Extended Data Fig. 2 | Marker enrichment at pioneered enhancers.** Enrichments (mean ± standard error of the mean) from ChIP-qPCR of the indicated markers or ATAC-qPCR at no (blue) or 72 h (red) of Pax7 activation. Enrichments are expressed relative to mean enrichment at random loci (Pax7 ChIP) or constitutive enhancers (all others). In each case, data is shown for one of duplicate or triplicate experiments at the panel of 14 pioneered enhancers (Pio) as well as at constitutive enhancers (Con) and random loci (Rnd). Right boxplots represent mean enrichments at the 14 pioneered enhancers and the p-value (two-tailed paired t-test) compares enrichments at no vs. 72 h Pax7 activation.

**Extended Data Fig. 3 | Variations in Pax7 pioneering kinetics and initial chromatin state. a, b,** Mean (± standard error of the mean, n ≥ 2) ChIP/ATAC-qPCR time courses of the indicated markers at the panel of 14 pioneered enhancers split to the stronger and the weaker third subsets according to Pax7 recruitment at 48 h. The sequence of events is preserved but a delay of about 6 h in markers $t_{1/2}$ is observed for the weaker sites compared to the stronger ones. **c,** H3K4me1 ChIP-Seq time courses at all genome-wide pioneered enhancers. Sites color-coded according to Pax7 recruitment at 1 h and 48 h. **d,** Correlations between the indicated parameters at the 14 pioneered enhancers. Significant correlations are observed between the initial H1 levels, the initial Pax7 recruitment (1 h) up to 48 h and the overall speed of the pioneering process. Theses correlations are lost for long Pax7 activation (> 20 days) or when stably expressed suggesting that pioneered enhancers ultimately reach a similar final state. No correlation is observed with the initial levels of H3K9me2 or total histone H3. P-values from two-tailed t-tests (*** ≤ 0.001, ** ≤ 0.01, * ≤ 0.5, – otherwise).

**Extended Data Fig. 4 | Sequential Pax7 recruitment and chromatin opening at the 14 pioneered enhancers.** Pax7 ChIP-Seq and ATAC-Seq profiles at the panel of 14 pioneered enhancers studied by qPCR at the indicated times after Tam addition and after stable expression.

**Extended Data Fig. 5 | Characterization of mimosine-arrested cells. a,** FACS profiles of AtT20 cells treated with mimosine. Cells arrested in G1 by 24 h mimosine treatment are viable and re-enter cell cycle following release from mimosine-induced cell cycle arrest (n = 2). **b,** p300, Brg1 and Tpit western blots in nuclear extracts of normal and cells treated for 60 h with mimosine. The treatment does not affect p300 or Brg1 levels but results in a two-fold decrease of Tpit expression. A representative (n = 3) blot is shown.

**Extended Data Fig. 6 | Pioneering assessment in mimosine-arrested AtT20 cells.** ChIP/ATAC-qPCR data for individual sites assessed in normal and mimosine-arrested cells following Pax7 activation with tamoxifen. One of two biological experiments is shown for each marker. P-values from two-tailed t-tests (*** ≤ 0.001, ** ≤ 0.01, * ≤ 0.5, – otherwise).

**Extended Data Fig. 7 | Time course of KDM3A recruitment and further characterization of Pax7-primed enhancers. a,** Mean (± standard error of the mean, n ≥ 2) ChIP-qPCR time courses of KDMA3A compared to Pax7 recruitment at the 14 pioneered enhancers. **b,** Heatmaps of ChIP-Seq for the indicated markers at multiple subsets of Pax7-targeted sites with or without stable Pax7 expression.

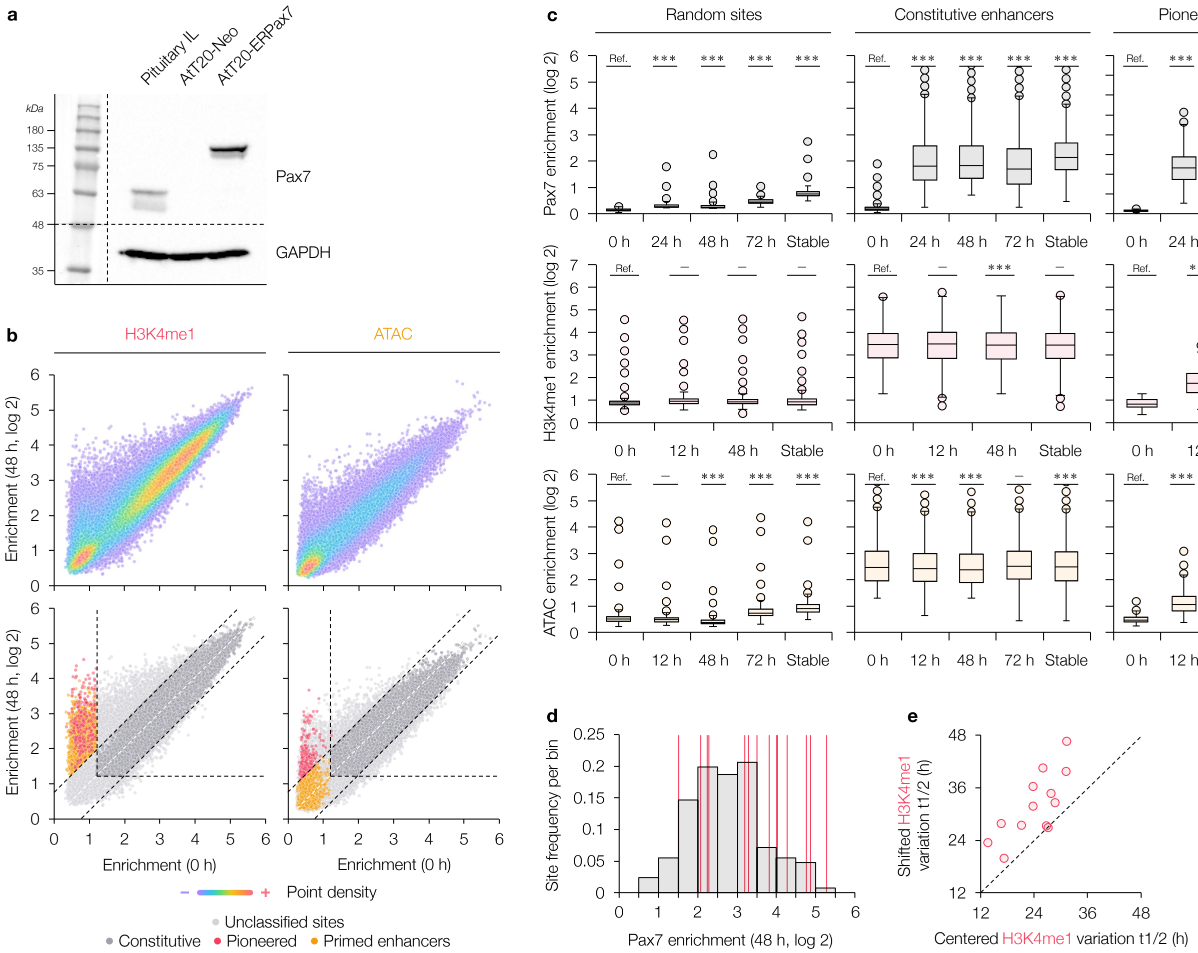


### Extended Data Fig. 1

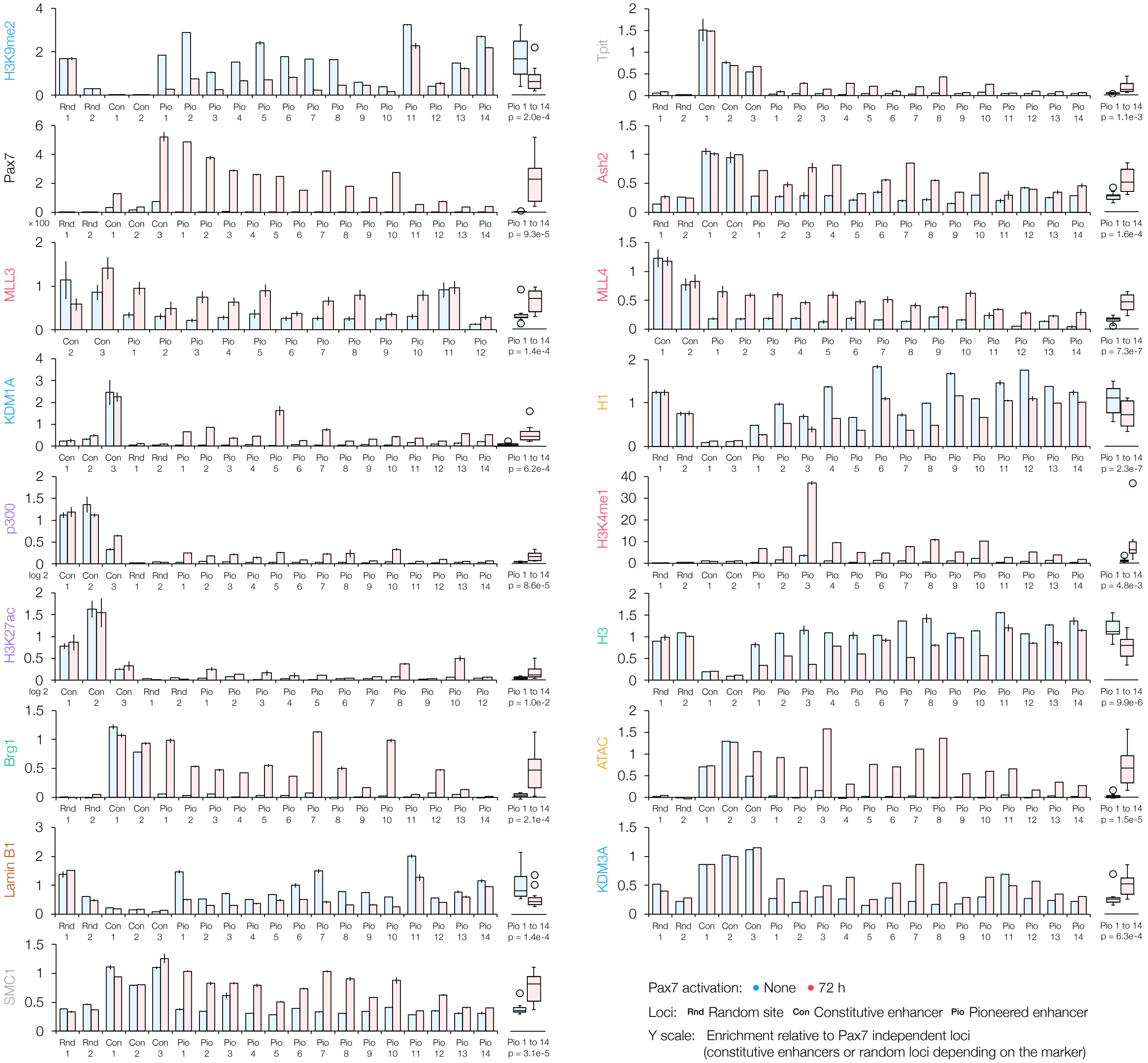


**Extended Data Fig. 2**

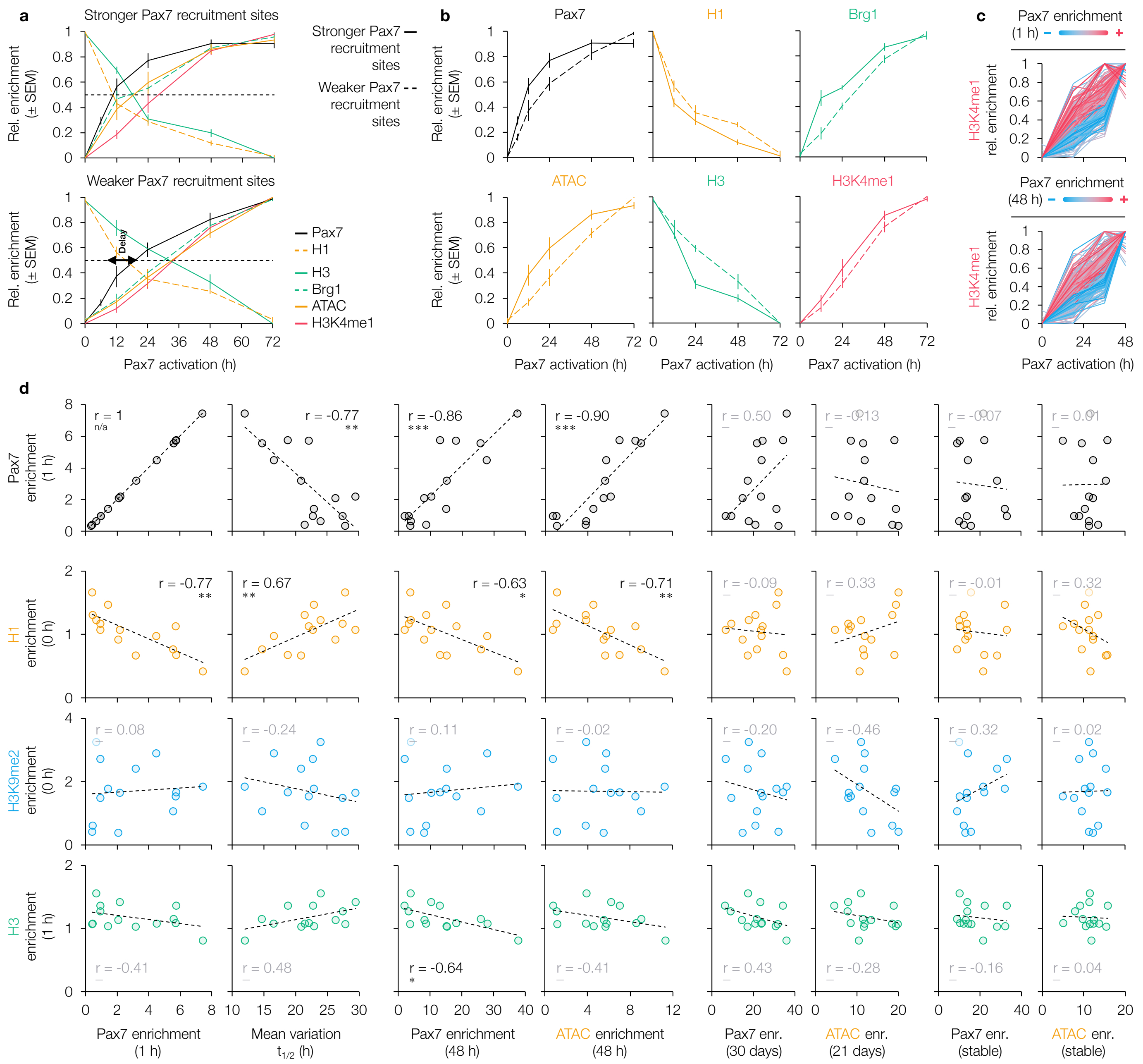


**Extended Data Fig. 3**

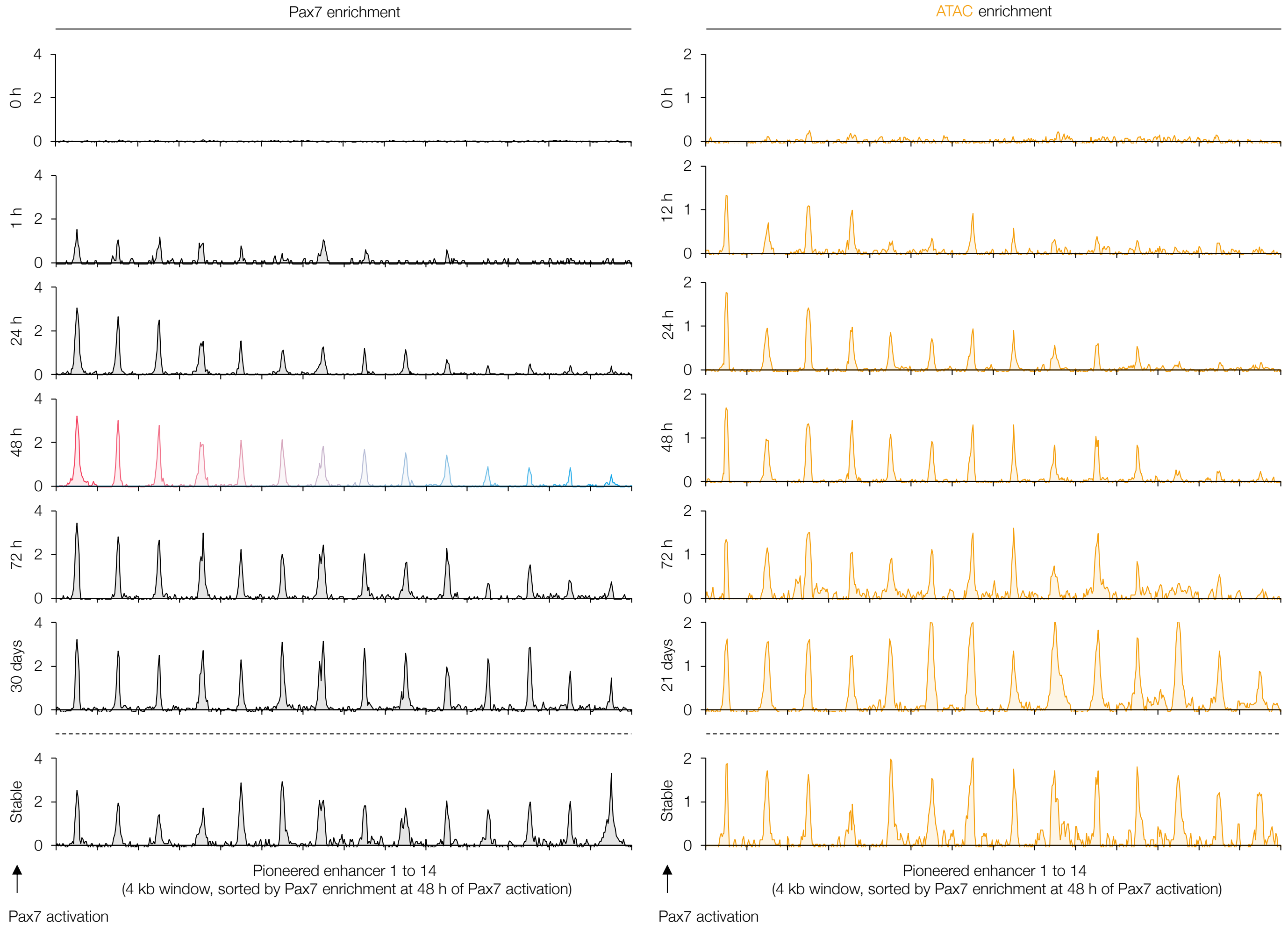


**Extended Data Fig. 4**

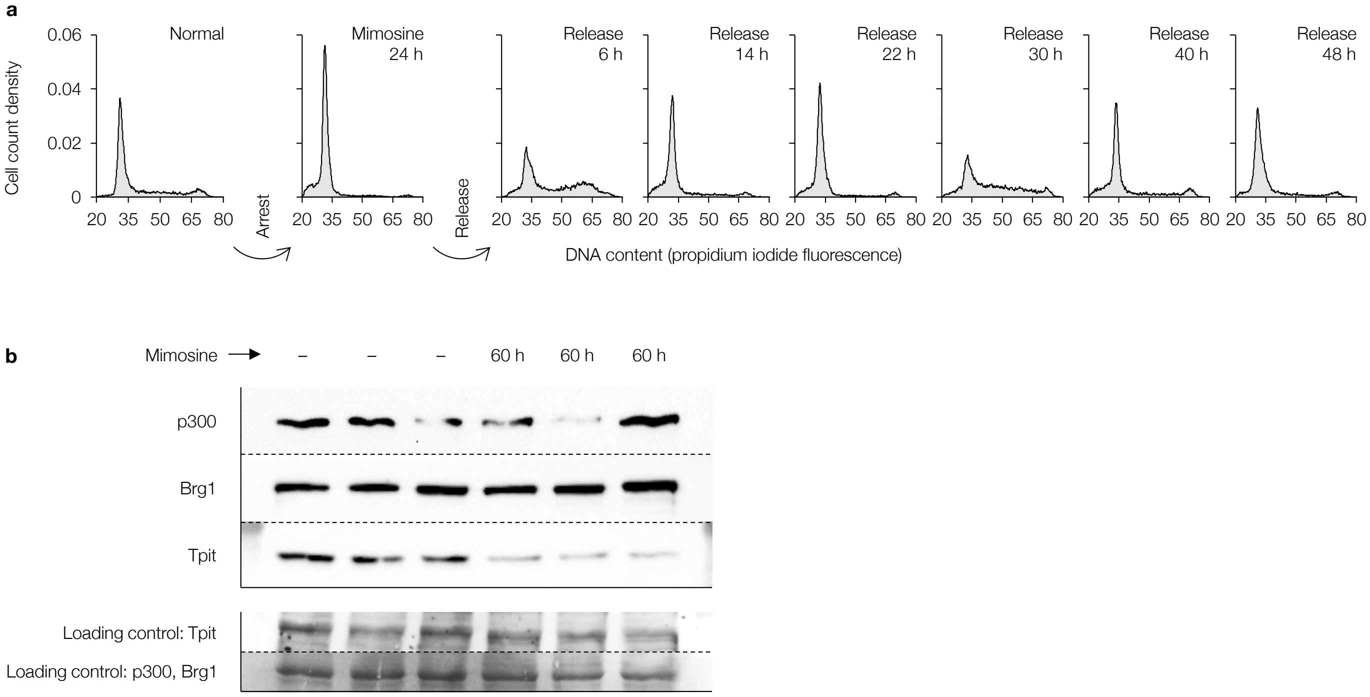


**Extended Data Fig. 5**

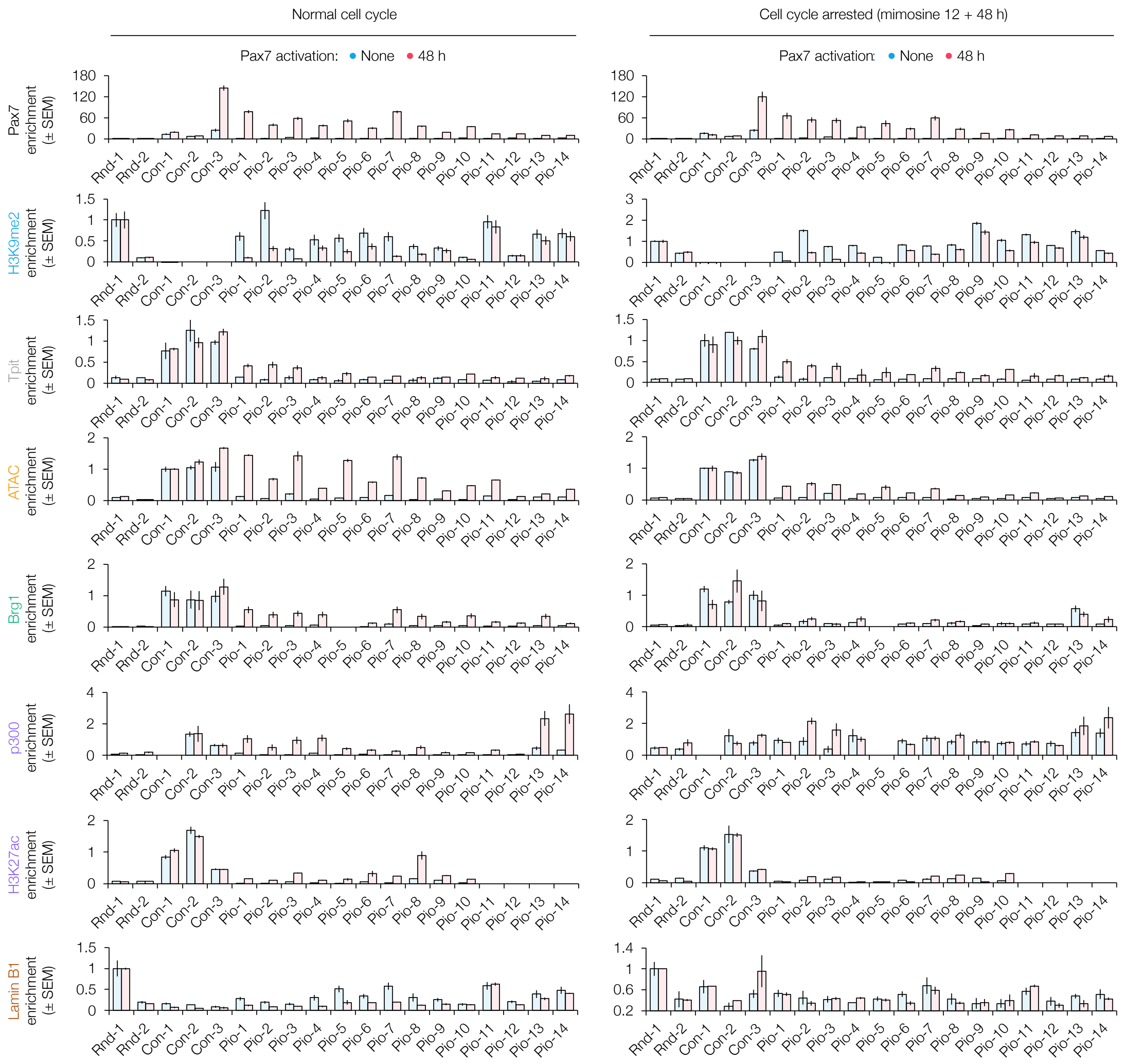


**Extended Data Fig. 6**

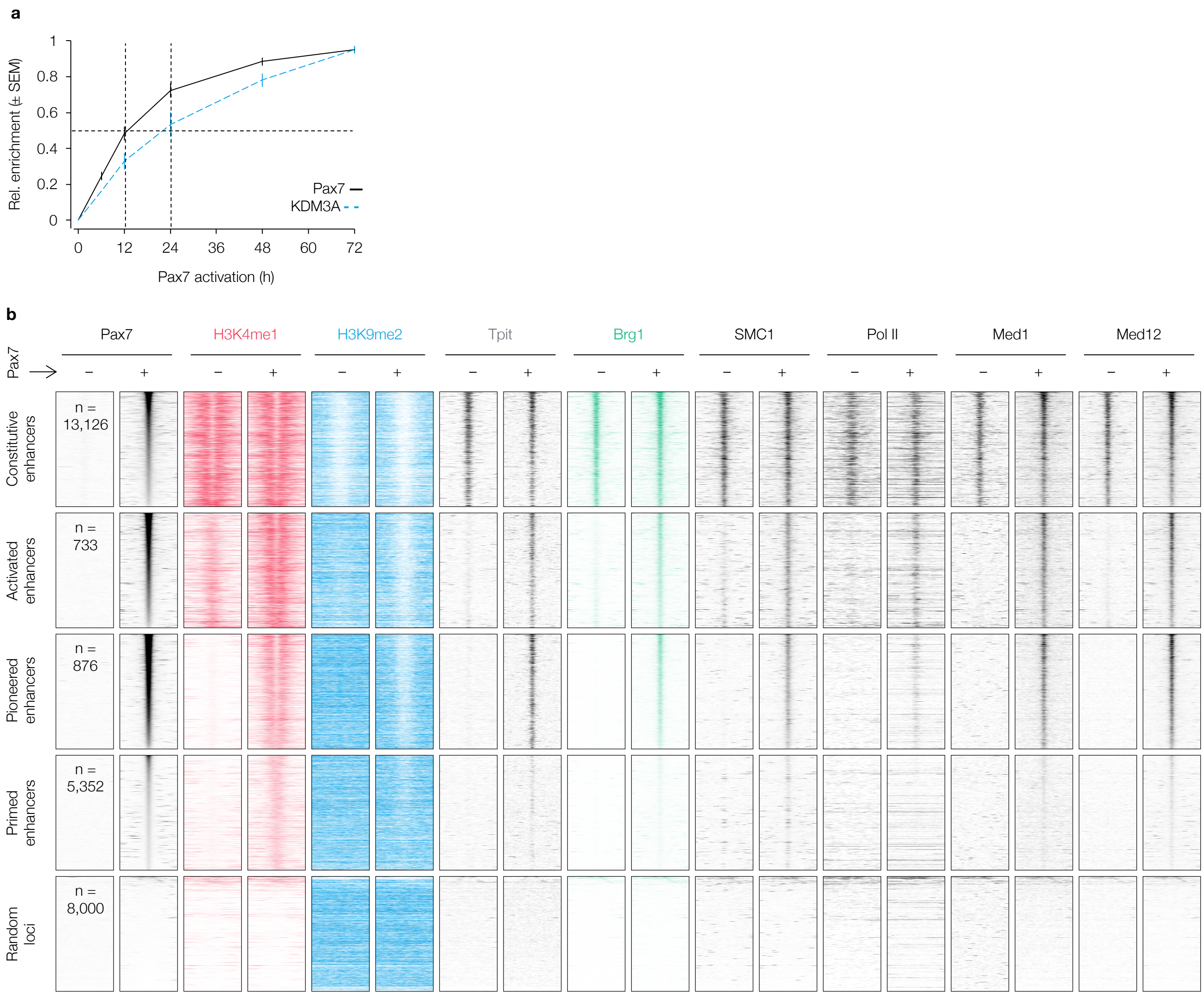


**Extended Data Fig. 7**
